# Fluorescent in situ sequencing of DNA barcoded antibodies

**DOI:** 10.1101/2020.04.27.060624

**Authors:** Richie E. Kohman, George M. Church

## Abstract

Biological tissues contain thousands of different proteins yet conventional antibody staining can only assay a few at a time because of the limited number of spectrally distinct fluorescent labels. The capacity to map the location of hundreds or thousands of proteins within a single sample would allow for an unprecedented investigation of the spatial proteome, and give insight into the development and function of diseased and healthy tissues. In order to achieve this goal, we propose a new technology that leverages established methodologies for in situ imaging of nucleic acids to achieve near limitless multiplexing. The exponential scaling power of DNA technologies ties multiplexing to the number of DNA nucleotides sequenced rather than the number of spectrally distinct labels. Here we demonstrate that barcode sequencing can be applied to in situ proteomics by sequencing DNA conjugated antibodies bound to biological samples. In addition, we show a signal amplification method which is compatible with barcoded antibodies.

## Main

The selective staining of proteins within cells and tissues is routinely performed in basic and applied research with application ranging from the diagnosis of patient samples to the characterization of fundamental, developmental processes. For decades, fluorescent antibodies have been used to probe the myriad of spatially distributed molecular targets within biological samples.^1^ Despite their widespread use, however, the number of targets that can be imaged is relatively small compared to the total diversity in biological samples, and it is often limited by the number of dye colors that can be easily distinguished. One approach to circumvent the color-space limit is iterative staining methods, which enable the re-use of colors for distinct targets by separating the labelling reaction into multiple cycles.^2^ Direct removal of antibodies,^3, 4^ and photobleaching of dyes^5^ have been demonstrated, however these strategies require harsh conditions and are damaging to the samples. In situ hybridization (ISH) of oligonucleotides conjugated to antibodies has emerged as a mild and reversible way to address a large number of targets with iterative staining.^6–9^ Although effective, ISH only scales linearly with the number of iterative labelling reactions and does not take advantage of the data density of DNA (4^N^ where N = the number of bases in the oligos). For instance, one 20 nucleotide-long ISH probe encodes 1 bit of information whereas the sequencing of this length encodes 4^20^ (over 1 trillion) bits. Recently, Goltsev *et al* have shown a sequencing based approach to read out the location and identity of barcoded antibodies^10^, however the chemistry chosen did not take advantage of high density base space and therefore drastically reduced the sequence diversity that could be investigated. Here we demonstrate in situ sequencing of oligonucleotide-conjugated antibodies bound to their biological targets to read out the spatial proteome. By directly using Sequencing by Synthesis (SBS) chemistry (**Supplemental Figure 1**)^11^ on stained tissue samples, we show that the unique barcode of an antibody can be sequenced in multiple imaging rounds and the nucleotide identity linked to antibody identity. We demonstrate our approach on cellular targets in whole mouse brain slices and additionally describe an amplification technique that can be used to increase the signal of our antibodies.

First we developed a straightforward method to conjugate oligonucleotides to antibodies using a variation of previous approaches (**Supplemental Figure 2**).^12^ In short, lysine residues on the antibodies were reacted with a commercially available linker to display DBCO groups. Azide-containing oligonucleotides were reacted with these groups via copper-free click chemistry at a stoichiometry that ensured no unmodified antibody remained, eliminating the possibility that they could outcompete the bulkier conjugated product for binding. Our protocol routinely produced an average of five oligos per antibody over antibodies raised in a variety of species.

Next we validated our antibody conjugates against fluorescently tagged versions to ensure binding was not altered by the presence of DNA. We chose to stain against glial fibrillary acidic protein (GFAP) in mouse brain slices because this protein is cell-filling in glial cells and therefore produces unambiguous, stellate-like cell images when staining is successful. For proof of concept we used a secondary antibody approach and visualized our barcoded antibodies with a fluorescent in situ hybridization probe (**Figure 1A**). When comparing our barcoded antibody against the commercially-available control, we discovered the staining quality was comparable. Examination of brain hemispheres indicated good staining throughout the whole section with GFAP positive cells clearly visible (**Figure 1B, C**). With respect to cell morphology, there was no observable difference between the results from the two antibodies. Quantification of the staining intensity showed that the signals between the two conditions were within an order of magnitude (with the barcoded antibody being approximately 60% as bright) indicating our degree of substitution of conjugation was useful for staining (**Supplemental Figure 3A**).

**Figure 1.**
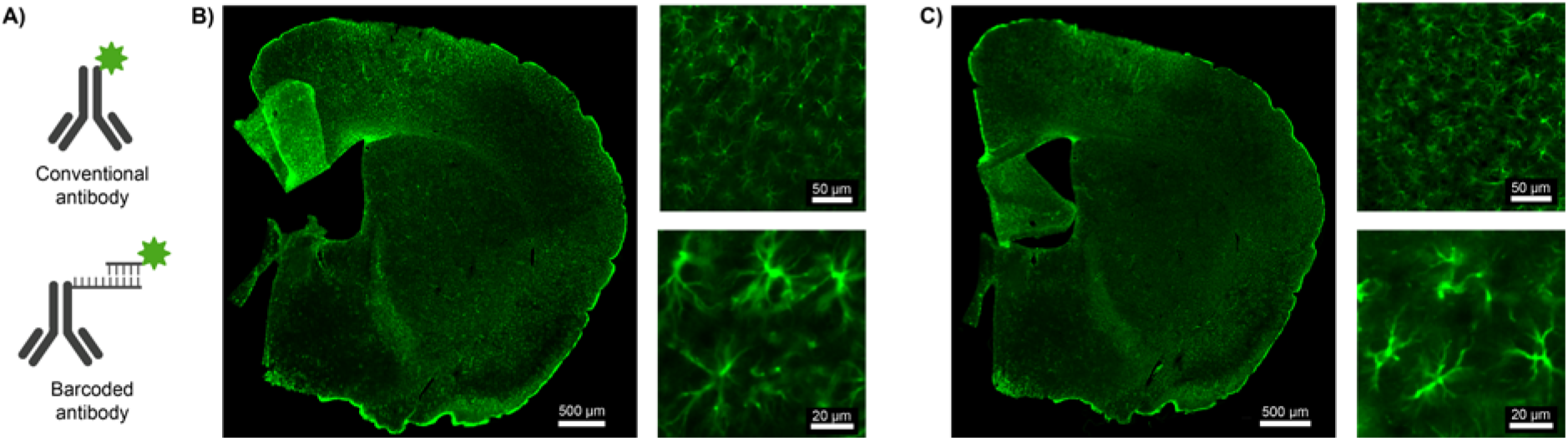
Staining of barcoded versus conventional antibodies against glial fibrillary acidic protein (GFAP) in mouse brain slices. A) Schematic depiction of antibodies. Conventional antibodies contain fluorescent dyes directly attached whereas barcoded antibodies contain oligonucleotides which can be imaged with a fluorescent complementary oligo. Images of a mouse brain hemisphere as well as different regions of interest stained with B) conventional or C) barcoded antibodies against GFAP. Observed cellular morphology was good for both conditions and staining intensity was comparable.

With validated antibodies in hand, we next evaluated the ability to sequence the barcoded antibodies in situ. We chose to use Illumina NextSeq SBS chemistry which encodes four bases with two easily distinguishable colors.^13^ This encoding is made possible by having two bases encoded by each color, one base encoded by two color and another encoded with no colors (**Figure 2A**). We included an orthogonal fluorophore at the 5’ end of our sequencing primer to provide a means for monitoring signal loss over sequencing steps. Stained slices were directly incubated with the sequencing reagents, washed, and imaged. We observed sustained fluorescence in our primer channel indicating that hybridization withstood the sequencing conditions and also fluorescent signal in our sequencing channels in excellent agreement with the predicted base (**Figure 2B**). For the first base bright signal in the green (561 nm) channel was observed in the same location as our primer signal and negligible signal was observed in the red channel. This color encodes the incorporation of adenine consistent to what was predicted. Importantly, signal was only observed in GFAP positive cells. Signal was not observed outside of the boundaries of the cells despite the potential that nicked genomic DNA present throughout the all of the cells in the tissue would serve as a source for sequencing priming. After imaging, a cleave solution was added that simultaneously removed the fluorophores and the 3’ OH blocking group. Cleavage caused the sequencing signals to disappear whereas the primer signal remained unchanged. Two additional iterations were then performed, both of which produced proper base calling and retention of primer signal. Both of these rounds produced dual colored bases as predicted. We observed clear signal in the GFAP positive cells but noticed a general increase in background staining, especially with the red (640 nm) channel (**Supplemental Figure 3B**). We expect the use of current tissue clearing techniques to be useful to eliminate this effect. Together these three rounds demonstrate proof of concept of in situ sequencing of barcoded antibodies and we expect multiplexing limits using this technique to be determined by the availability of antibodies rather than their readout.

**Figure 2.**
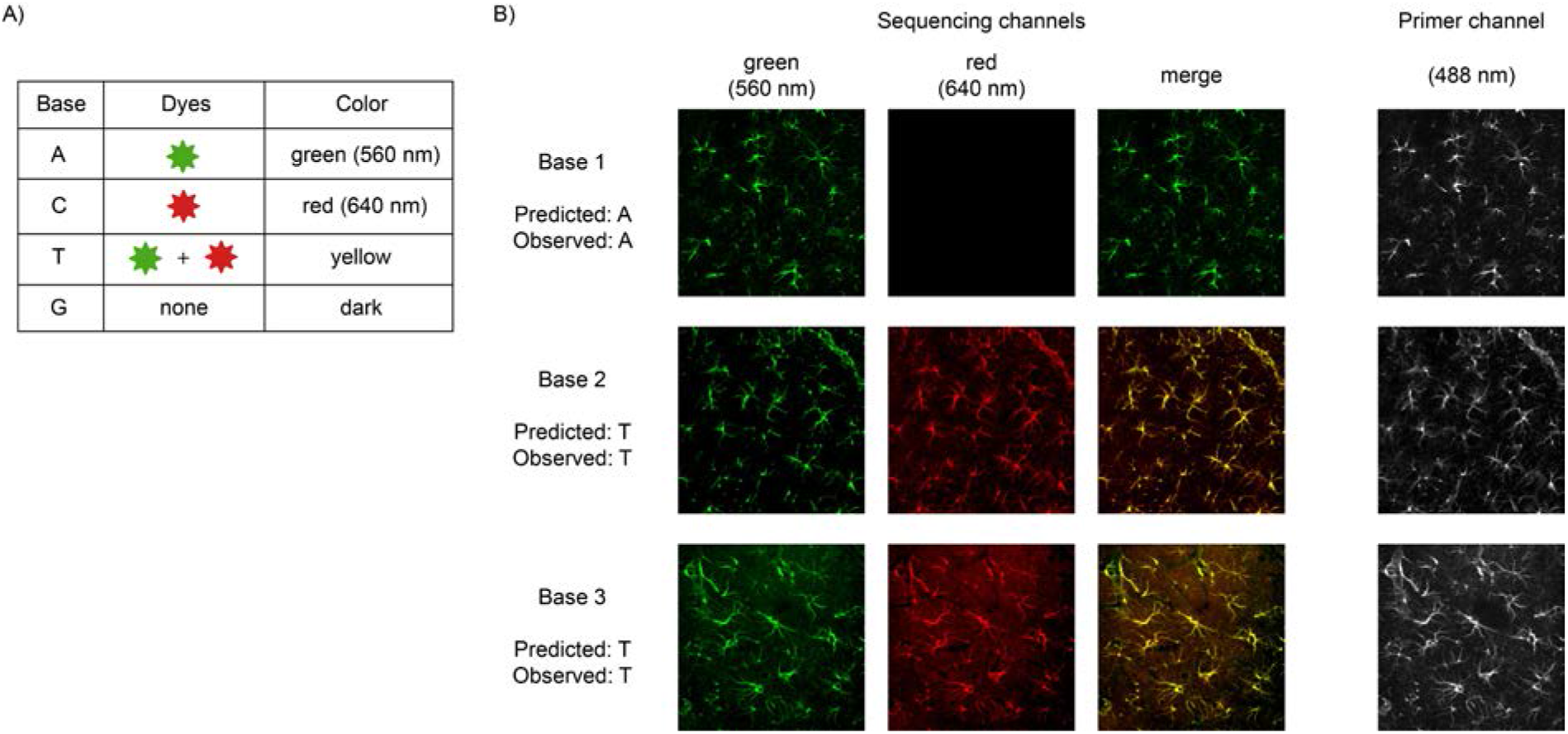
In situ sequencing of DNA barcoded antibodies. A) Base encoding scheme. Using two dyes, bases are encoded with individual dyes, the combination of both dyes, and no dyes. B) Three round of sequencing of GFAP barcoded antibodies. Individual and merged images show the correct color combinations for the predicted bases. Additionally, a separate dye on the sequencing primer confirms the position of the staining location.

Finally, we created a signal amplification technique that would enable the imaging of low-abundance targets. Several amplification techniques for barcoded antibodies have been developed by our group and others,^14–16^ however these require design criteria which limit their ability to scale to hundreds or thousands of targets. Here we chose to use a padlock probe approach which forms a circular template to our barcode and enables rolling circle amplification (RCA) after ligation.^17–19^ Previously we have shown the paralleled use of over 40,000 padlock probes in the ex situ context,^20^ and we and others have routinely used RCA for in situ RNA sequencing^21–24^. A similar approach called immuno-RCA^25, 26^ has been demonstrated ex vitro however it has not been applied for multiplexed in situ proteomics. We performed padlocking and RCA directly on tissue sections stained with our barcoded antibodies and noticed a strong reaction time dependency with respect to the signal quality. For GFAP samples, 20 minutes was insufficient to produce adequate signal, and punctate fluorescence was observed rather than well resolved cells (**Supplemental Figure 4A**). In contrast, 2.5 hours produced the desired stellate cell shapes indicative of proper staining (**Figure 3**). When compared with unamplified samples, staining signal was found to be approximately 5 times as intense (**Supplemental Figure 3C**). We also discovered that over-amplification could be problematic. When RCA was performed overnight, we observed extremely bright tissues slices that did not possess recognizable staining. Additionally these tissues became difficult to work with and had a tendency to irreversibly fold on top of themselves (**Supplemental Figure 4B**). This observation is in stark contrast to the amplicons observed from single molecule RNA detection which appear as bright, circular objects rather than large aggregates.^21–24^ This experiment highlights the ability to tune the signal intensity of a given target within a tailored time window.

**Figure 3.**
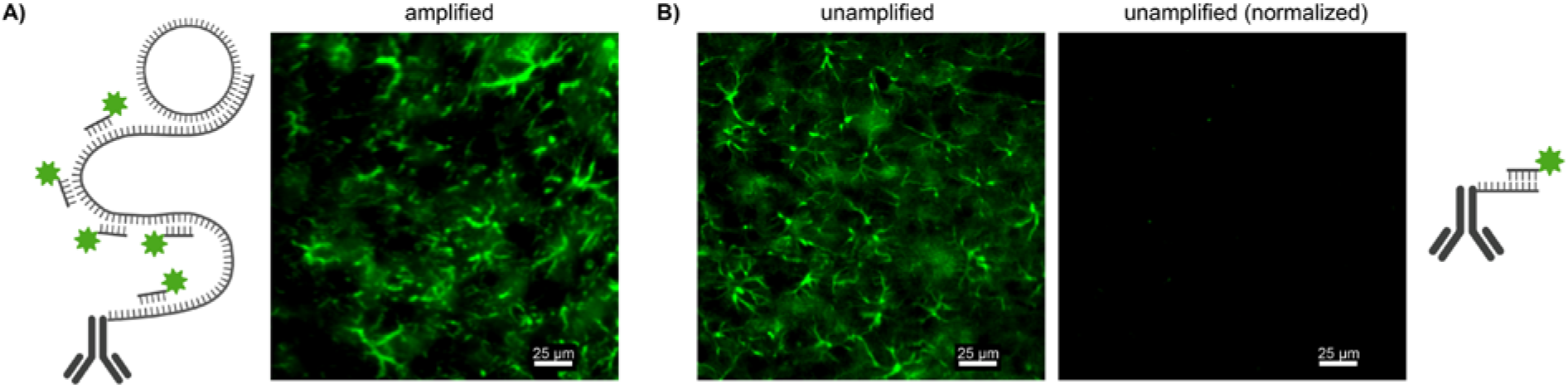
Amplification of barcoded antibody signal using rolling circle amplification (RCA). A) GFAP signal after 2.5 hours of RCA. B) Unamplified GFAP signal from barcoded antibodies. The left image shows adjusted intensities to display staining signals while the right image is normalized to A) to give a direct comparison of intensity.

In summary, we have demonstrated a technique to perform multiplexed spatial proteomics using DNA barcoded-antibodies in stained tissue samples. This approach is straightforward and should scale to numerous targets in parallel provided orthogonal antibodies are available. We demonstrated three rounds of in situ sequencing which corresponds to up to 64 unique targets which could potentially be assayed in a single experiment. For comparison, in situ hybridization of barcoded antibodies for three rounds using two colors would only be able to detect 6 unique targets. We expect straightforward scaling of our approach to large numbers of proteins through the use of barcoded primary antibodies.^14^ We also demonstrated a rolling circle amplification method which should allow sequencing of targets present in low amounts. Our technique should be compatible with in situ sequencing of other molecular types such as RNA or genomic DNA and we foresee the approach described here as a starting point for spatial multi-OMIC assays of cells and tissues.

## Acknowledgements

We thank Dr. Andrew Pawlowski for his suggestions on protocol details and David Kalish for help with image analysis. This work is supported by NIH grant RM1HG008525, NIH grant 1R01NS112716-01A1, IARPA grant D16PC0008 (MICrONS), and funding from the Wyss Institute for Biologically Inspired Engineering at Harvard.

## Author Contributions

REK and GMC conceived the project. REK designed and performed the experiments. REK and GMC wrote the manuscript.

## Competing Interests

REK was a founding scientist and has equity in Expansion Technologies and GMC is cofounder and has equity in ReadCoor and ManifoldBio.

## Notes

### Competing Interest Statement

REK was a founding scientist and has equity in Expansion Technologies and GMC is co-founder and has equity in ReadCoor and ManifoldBio.

## References

1. Buchwalow, I. B.; Böcker, W., Immunohistochemistry: basics and methods. Springer: Heidelberg; Berlin, 2010.

2. Dixon, A. R.; Bathany, C.; Tsuei, M.; White, J.; Barald, K. F.; Takayama, S., Recent developments in multiplexing techniques for immunohistochemistry. Expert Review of Molecular Diagnostics 2015, 15 (9), 1171–1186.

3. Micheva, K. D.; Smith, S. J., Array Tomography: A New Tool for Imaging the Molecular Architecture and Ultrastructure of Neural Circuits. Neuron 2007, 55 (1), 25–36.

4. Ku, T.; Swaney, J.; Park, J.-Y.; Albanese, A.; Murray, E.; Cho, J. H.; Park, Y-G.; Mangena, V.; Chen, J.; Chung, K., Multiplexed and scalable super-resolution imaging of three-dimensional protein localization in size-adjustable tissues. Nature Biotechnology 2016, 34 (9), 973–981.

5. Schubert, W.; Bonnekoh, B.; Pommer, A. J.; Philipsen, L.; Böckelmann, R.; Malykh, Y.; Gollnick, H.; Friedenberger, M.; Bode, M.; Dress, A. W. M., Analyzing proteome topology and function by automated multidimensional fluorescence microscopy. Nature Biotechnology 2006, 24 (10), 1270–1278.

6. Chen, F.; Tillberg, P. W.; Boyden, E. S., Expansion microscopy. Science 2015, 347 (6221), 543.

7. Wang, Y.; Woehrstein, J. B.; Donoghue, N.; Dai, M.; Avendaño, M. S.; Schackmann, R. C. J.; Zoeller, J. J.; Wang, S. S. H.; Tillberg, P. W.; Park, D.; Lapan, S. W.; Boyden, E. S.; Brugge, J. S.; Kaeser, P. S.; Church, G. M.; Agasti, S. S.; Jungmann, R.; Yin, P., Rapid Sequential in Situ Multiplexing with DNA Exchange Imaging in Neuronal Cells and Tissues. Nano Letters 2017, 17 (10), 6131–6139.

8. Agasti, S. S.; Wang, Y.; Schueder, F.; Sukumar, A.; Jungmann, R.; Yin, P., DNA-barcoded labeling probes for highly multiplexed Exchange-PAINT imaging. Chemical Science 2017, 8 (4), 3080–3091.

9. Kohman, R. E., Combining Protein Barcoding with Expansion Microscopy for in-situ, Spatially-resolved Proteomics. US Patent App. 15/446,005 2017.

10. Goltsev, Y.; Samusik, N.; Kennedy-Darling, J.; Bhate, S.; Hale, M.; Vazquez, G.; Black, S.; Nolan, G. P., Deep Profiling of Mouse Splenic Architecture with CODEX Multiplexed Imaging. Cell 2018, 174 (4), 968–981.e15.

11. Guo, J.; Yu, L.; Turro, N. J.; Ju, J., An Integrated System for DNA Sequencing by Synthesis Using Novel Nucleotide Analogues. Accounts of Chemical Research 2010, 43 (4), 551–563.

12. Gong, H.; Holcomb, I.; Ooi, A.; Wang, X.; Majonis, D.; Unger, M. A.; Ramakrishnan, R., Simple Method To Prepare Oligonucleotide-Conjugated Antibodies and Its Application in Multiplex Protein Detection in Single Cells. Bioconjugate Chemistry 2016, 27 (1), 217–225.

13. https://www.illumina.com/science/technology/next-generation-sequencing/sequencing-technology/2-channel-sbs.html.

14. Wang, Y.; Xie, W.; Kohman, R. E.; Church, G. M., Multiplexed imaging using same species primary antibodies with signal amplification. bioRxiv 2018.

15. Lin, R.; Feng, Q.; Li, P.; Zhou, P.; Wang, R.; Liu, Z.; Wang, Z.; Qi, X.; Tang, N.; Shao, F.; Luo, M., A hybridization-chain-reaction-based method for amplifying immunosignals. Nature Methods 2018, 15 (4), 275–278.

16. Saka, S. K.; Wang, Y.; Kishi, J. Y.; Zhu, A.; Zeng, Y.; Xie, W.; Kirli, K.; Yapp, C.; Cicconet, M.; Beliveau, B. J.; Lapan, S. W.; Yin, S.; Lin, M.; Boyden, E. S.; Kaeser, P. S.; Pihan, G.; Church, G. M.; Yin, P., Immuno-SABER enables highly multiplexed and amplified protein imaging in tissues. Nature Biotechnology 2019, 37 (9), 1080–1090.

17. Nilsson, M.; Dahl, F.; Larsson, C.; Gullberg, M.; Stenberg, J., Analyzing genes using closing and replicating circles. Trends in Biotechnology 2006, 24 (2), 83–88.

18. Gusev, Y.; Sparkowski, J.; Raghunathan, A.; Ferguson, H.; Montano, J.; Bogdan, N.; Schweitzer, B.; Wiltshire, S.; Kingsmore, S. F.; Maltzman, W.; Wheeler, V., Rolling Circle Amplification: A New Approach to Increase Sensitivity for Immunohistochemistry and Flow Cytometry. The American Journal of Pathology 2001, 159 (1), 63–69.

19. Kazane, S. A.; Sok, D.; Cho, E. H.; Uson, M. L.; Kuhn, P.; Schultz, P. G.; Smider, V. V., Site-specific DNA-antibody conjugates for specific and sensitive immuno-PCR. Proceedings of the National Academy of Sciences 2012, 109 (10), 3731.

20. Li, J. B.; Levanon, E. Y.; Yoon, J.-K.; Aach, J.; Xie, B.; LeProust, E.; Zhang, K.; Gao, Y.; Church, G. M., Genome-Wide Identification of Human RNA Editing Sites by Parallel DNA Capturing and Sequencing. Science 2009, 324 (5931), 1210.

21. Lee, J. H.; Daugharthy, E. R.; Scheiman, J.; Kalhor, R.; Yang, J. L.; Ferrante, T. C.; Terry, R.; Jeanty, S. S. F.; Li, C.; Amamoto, R.; Peters, D. T.; Turczyk, B. M.; Marblestone, A. H.; Inverso, S. A.; Bernard, A.; Mali, P.; Rios, X.; Aach, J.; Church, G. M., Highly Multiplexed Subcellular RNA Sequencing in Situ. Science 2014, 343 (6177), 1360.

22. Lee, J. H.; Daugharthy, E. R.; Scheiman, J.; Kalhor, R.; Ferrante, T. C.; Terry, R.; Turczyk, B. M.; Yang, J. L.; Lee, H. S.; Aach, J.; Zhang, K.; Church, G. M., Fluorescent in situ sequencing (FISSEQ) of RNA for gene expression profiling in intact cells and tissues. Nature Protocols 2015, 10 (3), 442–458.

23. Chen, X.; Sun, Y.-C.; Church, G. M.; Lee, J. H.; Zador, A. M., Efficient in situ barcode sequencing using padlock probe-based BaristaSeq. Nucleic Acids Research 2017, 46 (4), e22–e22.

24. Iyer, E. P. R.; Punthambaker, S.; Liu, S.; Jindal, K.; Farrell, M.; Murn, J.; Ferrante, T.; Rudnicki, S.; Kohman, R. E.; Wassie, A. T.; Goodwin, D.; Chen, F.; Alon, S.; Sinha, A.; Milanova, D.; Aron, L.; Camplisson, C.; Skrynnyk, A.; Reginato, P. L.; Conway, N.; Aach, J.; Yankner, B.; Boyden, E. S.; Church, G. M., Barcoded oligonucleotides ligated on RNA amplified for multiplex and parallel in-situ analyses. bioRxiv 2018, 281121.

25. Schweitzer, B.; Wiltshire, S.; Lambert, J.; Malley, S.; Kukanskis, K.; Zhu, Z.; Kingsmore, S. F.; Lizardi, P. M.; Ward, D. C., Immunoassays with rolling circle DNA amplification: A versatile platform for ultrasensitive antigen detection. Proceedings of the National Academy of Sciences 2000, 97 (18), 10113.

26. Ali, M. M.; Li, F.; Zhang, Z.; Zhang, K.; Kang, D.-K.; Ankrum, J. A.; Le, X. C.; Zhao, W., Rolling circle amplification: a versatile tool for chemical biology, materials science and medicine. Chemical Society Reviews 2014, 43 (10), 3324–3341.

