## Supplemental Information for "Fluorescent in situ sequencing of DNA barcoded antibodies"

### Methods

#### Antibody-DNA Conjugation

To a solution of 100 µL (0.813 nmol) antibody (donkey anti mouse, Jackson ImmunoResearch, 715-005-151) was added 2.00 µL (20.0 nmol) of a 10 mM solution of DBCO-PEG13-NHS in DMSO (Aldrich). After 2 hours, the reaction was purified with a 50k Amicon spin filter (3 spins with PBS, 5 min & 14k x g per spin). The degree of functionalization was calculated by dividing the concentration of dibenzylcyclooctyne (DBCO) by the concentration of the antibody. Concentrations were calculated using Beer's law with absorbances measured using UV-vis spectroscopy (Nanodrop). Extinction coefficients of 204,000 $M^{-1}$ $cm^{-1}$ (at 280 nm) and 12,000 $M^{-1}$ $cm^{-1}$ (at 309 nm) were used for DBCO and antibody respectively. 15 µL of the antibody solution (0.359 nmol) was then reacted overnight with 32 µL (10 uM TE, 3.2 nmol) azido oligo used (100 µM). The final product was obtained at a similar concentration to the starting antibody and used without any further purification. This protocol gave comparable results using antibodies against mouse, rabbit, goat, sheep, chicken, guinea pig, and Armenian hamster (Jackson ImmunoResearch).

#### Tissue Acquisition

Tissue for this work was harvested from adult female wild-type C57BL/6J mice. All procedures were in accordance with the NIH Guide for Laboratory Animals and approved by the Harvard University Animal Care and Use and Biosafety Committees. Briefly, mice were anesthetized until unresponsive. Transcardial perfusion was performed first with PBS followed by 4% formaldehyde in PBS. The brain was removed and soaked in 4% formaldehyde in PBS overnight at 4ºC. The tissue was then transferred to a 30% sucrose in PBS solution. The brain was soaked for approximately 24 hours at 4ºC with mild mixing until it sank to the bottom of the container. It was cut in half down the midline and then embedded into OCT using a hexanes/dry ice bath before being cryosectioned at -20ºC at a thickness of 30 µm. Samples were collected free floating in PBS in 48 well plates. The solution was swapped to 70% ethanol in PBS for storage at 4 ºC until use.

#### Staining

All staining steps were performed with 200 µL of solution to samples in 48 well plates. Staining was performed using MAX Stain buffers (Active Motif). Slices were washed three times with MAXwash buffer and then incubated in MAXblock buffer for one hour with shaking. Anti-GFAP primary antibody (Cell Signaling Technologies, mouse mAb #3670) was then added at 1:100 in MAXstain buffer and incubated overnight at 4ºC with shaking. Slices were washed three times with MAXwash and then stained with secondary antibodies. For control staining, AlexaFluor488 modified antibody (donkey anti mouse, Jackson ImmunoResearch, 715-545-151) was added to slices at 1:500 dilution in MAXbind for approximately five hours before being washed with MAXwash. For sample staining, DNA barcoded antibody was added to slices at 1:100 dilution in crowing buffer for approximately five hours before being washed first with crowding buffer and then PBS. To visualize slices stained with barcoded antibodies, they were first washed with hybridization buffer (2x SSC, 10% formamide) and then incubated with “Barcode in situ hybridization probe/sequencing primer” at 500 nM in hybridization buffer for one hour. Slices were then wash with hybridization buffer and PBS prior to imaging.

Crowding buffer:
500 µL 20X SSC
500 mg 40kDa dextran (Aldrich)
5 µL TritonX (Aldrich)
3995 µL Pierce Protein-Free (TBS) Blocking Buffer (Thermo)

**Microscopy**
Widefield imaging was performed with a Nikon upright microscope using a 10X CFI Plan Apo Lambda objective lens (MRD00105, 0.45 NA) and FITC, Cy3, and C5 filter cubes. Confocal imaging was performed using a Nikon Ti inverted microscope containing a Yokogawa CSU-W1 spinning disk confocal head with an ORCA-Flash4.0 V3 Digital CMOS camera. Confocal imaging used a 20X water immersion CFI Apo LWD Lambda S objective lens (MRD77200, 0.95 NA) and 488, 561, and 640 nm laser lines.

**Sequencing**
Reagents for sequencing were utilized from Illumina NextSeq 500 Mid Output Kits V2 (FC-404-2003) cartridges. We refer to the contents of the largest buffer reservoir as Incorporation Buffer, the purple colored solution as Incorporation Mix, and the solution used to remove fluorescence as Cleave Solution. All reactions were performed in the free floating state using 200 µL of solution per sample. Sequencing began after addition of the “Barcode in situ hybridization probe/sequencing primer”. Slices were washed three times with Incorporation Buffer. Then a solution of 50% Incorporation Mix in Incorporation Buffer was added to the slices and incubated at 50ºC for 15 minute. This step was performed twice each time with fresh solution. Post-incorporation, the slices were washed with Incorporation Buffer at 50ºC for 15 minutes 3 times before imaging. After imaging, slices were incubated with Cleave Solution for 20 minutes at 50ºC. Samples were washed with Incorporation Buffer at 50ºC for 15 minutes 3 times before proceeding to the next round of sequencing.

**In situ amplification**
All steps were performed with 200 µL of solution to samples in 48 well plates. Amplification was performed on samples containing barcoded antibodies in the absence of a hybridized primer/imaging probe. Samples were first washed with hybridization buffer and then incubated with “Barcode padlock” at 100 nM in hybridization buffer at 37ºC

overnight. Next they were washed with hybridization buffer, PBS, and 1x Ligase Buffer. Samples were then incubated with ligation solution for 1 hours at 37ºC and 1 hour at 45ºC.

Ligation solution:
20 µL 10X Ligase Buffer
4 µL Taq DNA Ligase (NEB, 40U/µL)
10 µL 1M KCl
10 µL Formamide
156 µL Water

Samples were then washed with PBS and incubated with rolling circle amplification (RCA) solution for a designated period of time.

RCA solution:
20 µl phi29 (10U/µL)
20 µL 10X phi29 buffer
2 µL 20mM dNTP
2 µL BSA (20ug/µL)
20 µL 50% glycerol
1 µL 4mM aadUTP
135 µL H2O

Samples were then washed with PBS and incubated with $BS(PEG)_9$ solution for 1 hours to crosslink the aminoallyl-dUTP (aadUTP) groups in the amplicon. Samples were washed with 1M Tris-HCl 8.0 for 30 minutes and then with PBS. “Amplicon in situ hybridization probe” at 500 nM in hybridization buffer was added for one hour and then washed with hybridization buffer and PBS prior to imaging.

$BS(PEG)_9$ solution = 40 µL stock solution + 160 µL PBS
Stock solution = 100 mg $BS(PEG)_9$ + 465 µL DMSO

**Image analysis**

Analysis was performed using ImageJ/Fiji. For comparisons of antibody staining intensities, images were cropped to give areas of the same size in anatomically similar areas. In all cases, the edges of the brain and bright blood vessels were excluded to make the data representative of cellular staining. Cellular staining intensities were then calculated by taking the average intensity of the raw image data that resided within a mask. Masks were created by performing rolling ball background subtraction followed by thresholding. For the analysis of the sequencing data, a similar process was performed however multicolor images were first registered using the Linear Stack Alignment with SIFT plugin. For all cases, the mask was created using the data obtained from the fluorescent primer channel and applied to the sequencing channels.

**Oligo sequences (IDT)**

<u>Antibody barcode</u>
/5AzideN/TTTTCTAGCGTTAGGGTTAGCGAAGCGTTATAGCAGCGACTACGGT

Barcode in situ hybridization probe/sequencing primer
/5Alex488N/ACCGTAGTCGCTGCTATAACGCTTCGCTAACCC

Barcode Padlock
/5Phos/ACGCTTCGCTAACCCTACATGAGTCGTGAGTACGTTCGGATTCTTAGGGCGTAAGCCCGACCTATCTTCTTTACCGTAGTCGCTGCTATA

Amplicon in situ hybridization probe
GTTCGGATTCTTAGGGCGTA/3AlexF488N/

### Supplemental Figures

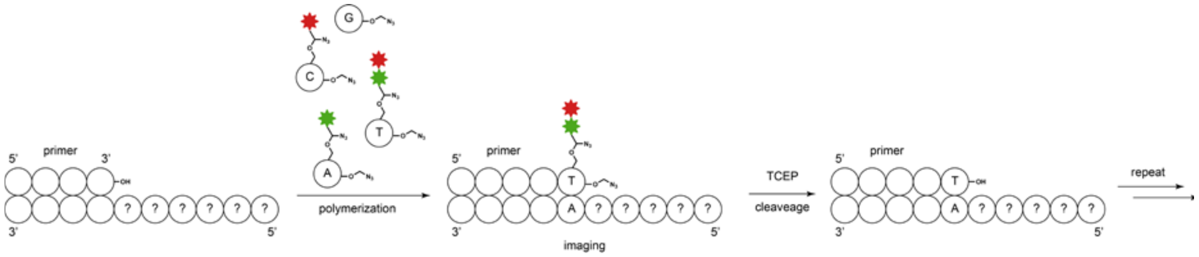


**Supplemental Figure 1.** Schematic depiction of Sequencing by Synthesis (SBS) chemistry. A polymerization step is performed off of a sequencing primer using a polymerase and 3’ blocked nucleoside triphosphates. The blocking group ensure only one base can be added and the color of the attached fluorophores uncovers the identity of the base. A cleavage step removes the fluorophore as well as the 3’ blocking group allowing the next sequencing iteration to be performed. Iterations are performed until the desired sequence is discovered.

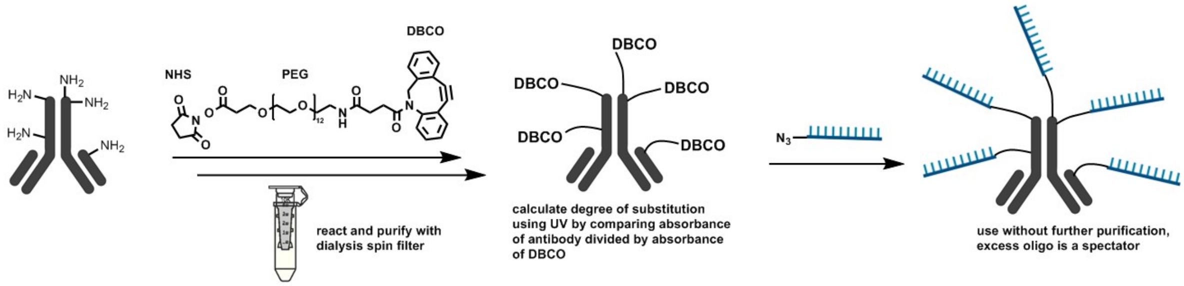


**Supplemental Figure 2.** Schematic depiction of oligonucleotide conjugation to antibodies. Oligos were attached in two steps. Purification was performed after the first step and degree of conjugation was obtained. Purchased azide-oligos were added in excess and not purified from the final product mixture so that accurate antibody concentration could easily be calculated by only factoring dilution.

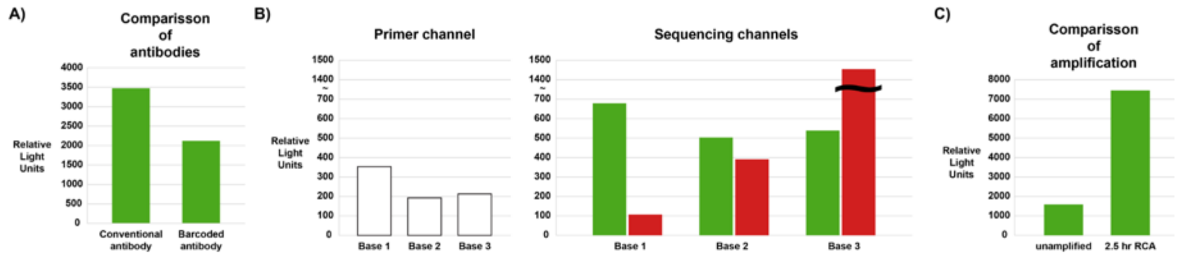


**Supplemental Figure 3.** Data graphs from tissue staining and sequencing. A) Comparison between the staining of GFAP in tissue using dye conjugated antibodies and oligonucleotide conjugated antibodies. B) Quantitative data from three rounds of sequencing. The left graph shows signal of the dye attached to the primer which changes little over the three rounds. The right graph shows the sequencing dyes which display base-specific intensities. Accumulation of the background staining was evident from the high intensity in the red channel at the third iteration. C) Comparison between staining with and without RCA amplification for 2.5 hours.

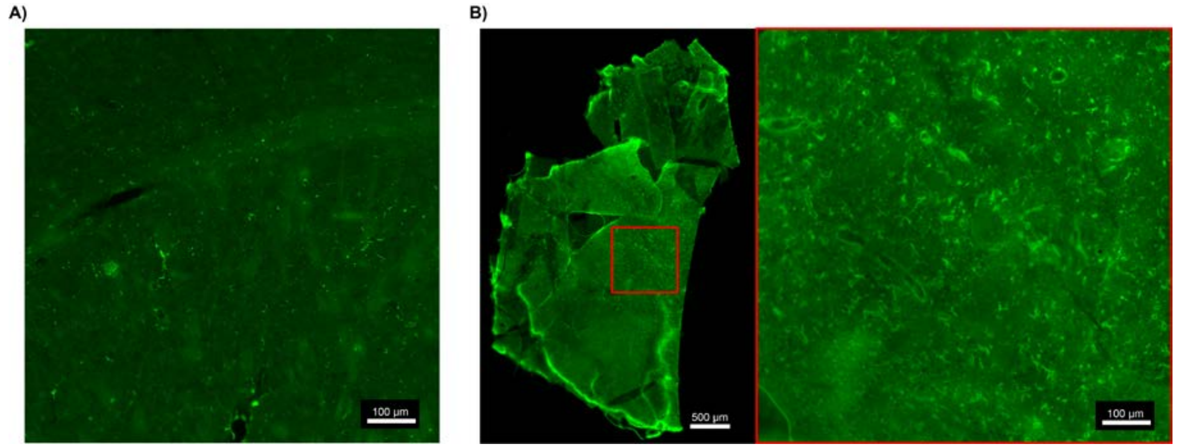


**Supplemental Figure 4.** Effect of amplification time on signal outcome. A) 20 minutes of RCA does not produce appreciable signal in the shape of GFAP-positive cells. B) Overnight RCA produces a tissue that is difficult to work with and has high background staining.
